# Hybrid machine-learning framework for volumetric segmentation and quantification of vacuoles in individual unlabeled yeast cells using holotomography

**DOI:** 10.1101/2023.06.18.545499

**Authors:** Moosung Lee, Marina Kunzi, Gabriel Neurohr, Sung Sik Lee, Yongkeun Park

**Author notes:** Contributed equally (Moosung Lee, Marina Kunzi).

## Abstract

The precise, quantitative evaluation of intracellular organelles in three-dimensional (3D) imaging data poses a significant challenge due to the inherent constraints of traditional microscopy techniques, the requirements of the use of exogenous labeling agents, and existing computational methods. To counter these challenges, we present a hybrid machine-learning framework exploiting correlative imaging of 3D quantitative phase imaging with 3D fluorescence imaging of labeled cells. The algorithm, which synergistically integrates a random-forest classifier with a deep neural network, is trained using the correlative imaging data set, and the trained network is then applied to 3D quantitative phase imaging of unlabeled cell data. We applied this method to unlabeled live budding yeast cells. The results revealed precise segmentation of vacuoles inside individual yeast cells, and also provided quantitative evaluations of biophysical parameters, including volumes, concentration, and dry masses of automatically segmented vacuoles.

## 1. Introduction

Biological cells, with their complex structure and specialized organelles separate from the cytoplasm, are the subject of increasingly intricate study. As the physical properties of intracellular compartments such as density and crowding come under focus [1, 2], their tightly regulated, functionally relevant characteristics have been revealed [3]. Deviations from normal density have been linked with altered cell function during starvation [4], cellular senescence [5], and differentiation [3, 6-8]. Furthermore, shifts in cytoplasm density can influence essential processes like phase separation [9-11], signaling [12, 13], and growth [14, 15]. However, the limited availability and scalability of methods for studying cell density at subcellular resolution pose significant challenges [16].

For instance, in budding yeast (*Saccharomyces Cerevisiae*), whole-cell density measurements are heavily skewed by the vacuole, an organelle with a function analogous to the mammalian lysosome and less dense than the rest of the cell. Therefore, an accessible, high-throughput method for automatic segmentation of vacuoles and cytoplasm is required.

Fluorescence (FL) microscopy, which marks the vacuole by tagging Vph1p with green fluorescent protein (GFP), has been instrumental in imaging intracellular compartments in budding yeast (Fig. 1a) [17]. Traditional 3D FL imaging methods entail stacking several axial images and implementing deconvolution algorithms. Alternatively, confocal or light sheet microscopy techniques can be employed for three-dimensional imaging acquisition. While these methods are effective in identifying intracellular structures, they may induce phototoxicity and provide limited information on the properties of the cytoplasm.

**Fig. 1.**
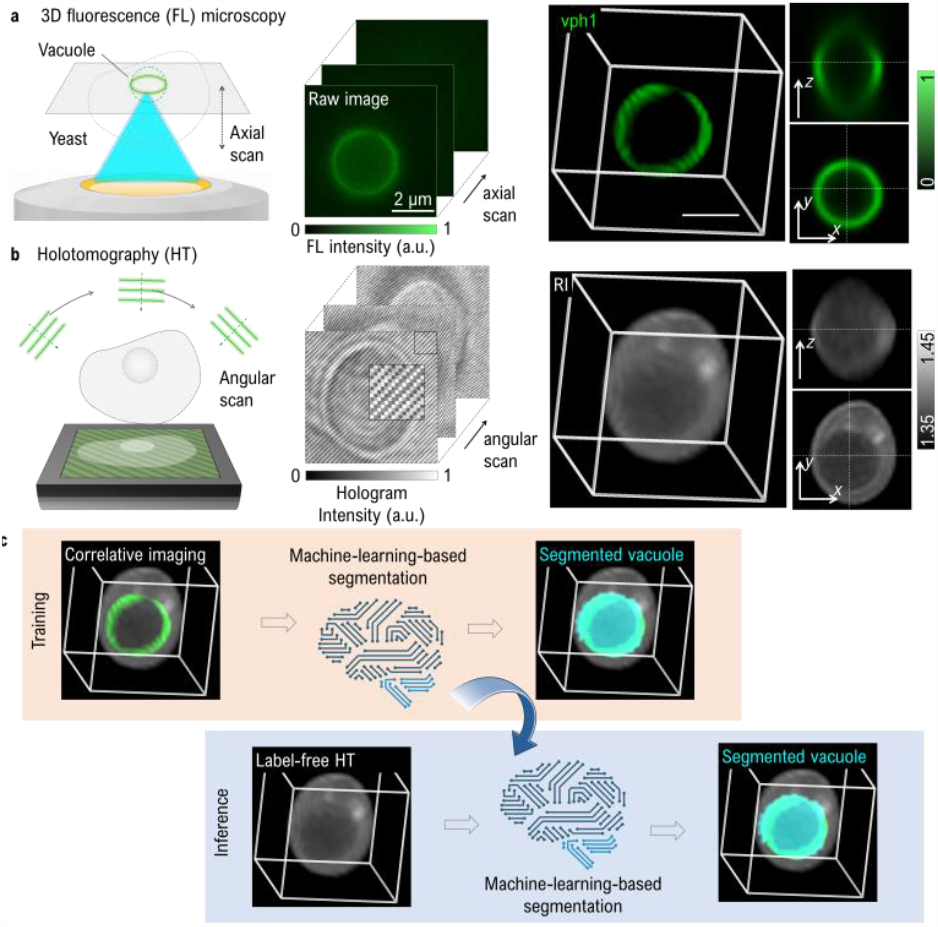
Volumetric segmentation via FL and RI correlative imaging. (a) Traditional FL microscopy utilizes axial scanning to generate a 3D image of Vph1-GFP. Although the deconvolved 3D data facilitate the identification of vacuole morphology, this method is constrained by photobleaching and limitations in quantifying cell density. (b) HT applies off-axis holography and angular scanning of incident plane waves to reconstruct an RI tomogram. While RI offers label-free contrast convertible to cell density, it lacks the required specificity of the vacuolar protein Vph1 for accurate analysis. (c) In this study, we employed correlative imaging to quantify cell density of vacuoles and cytoplasm with enhanced specificity. We also utilized machine learning for the automatic 3D segmentation of vacuoles.

Quantitative phase imaging (QPI) techniques offer a solution by providing a label-free method for precise quantification of intracellular organization [18] (Fig. 1b). These techniques employ interferometric microscopy to scan a plane wave angularly and reconstruct the volumetric distribution of the refractive index (RI). As RI increases proportionally with dry-mass concentration [19], this method offers dry-mass density as image contrast without the use of fluorescent labels. However, the lack of chemical specificity in RI imaging presents challenges for accurate segmentation of intracellular components.

To surmount these limitations, we have developed a hybrid machine-learning framework that utilizes correlative imaging of 3D QPI, also known as holotomography (HT), and 3D FL in labeled cells (Fig. 1c). The algorithm is trained using the correlative imaging dataset and subsequently applied to HT of unlabeled cell data. This method successfully merges the chemical specificity of fluorescence imaging with the quantitative capabilities of HT. Additionally, the incorporation of a machine learning framework enables automatic volumetric segmentation from raw images, thereby streamlining high-content analysis. Application of this method enabled us to quantitatively analyze the cytoplasm and vacuoles in unlabeled budding yeast cells. This study highlights its potential applicability in investigating intracellular density and its relationship with cellular function.

## 2. Methods

### 2.1 Sample preparation

Budding yeast cultures (*Saccharomyces Cerevisiae*, S288C BY4741) were cultured in synthetic complete (SC) medium at 30°C. Overnight grown cultures were diluted to an optical density (OD600) of 0.1 with fresh SC and allowed a 2-hour recovery period before imaging during the log phase or treating with rapamycin (0.1 or 1 μM) dissolved in ethanol. We imaged rapamycin-treated cells post 2-hour treatment. Stationary phase cells were imaged 24 hours after initial inoculation. For imaging purposes, we coated the glass bottom of the Tomodish (Tomocube Inc.) with 2 mg/mL Concanavalin A (Sigma) for 15 minutes, rinsed twice with SC, allowed the cells to attach for 5 minutes, followed by a single SC wash, and finally covered with a coverslip.

### 2.2 Optical imaging and reconstruction

For imaging budding yeasts, we used a HT system equipped with 3D FL imaging capability (HT-2H; Tomocube Inc.) [20]. The imaging system was equipped with water-immersion objective lenses (UPLASAPO60XW, numerical aperture = 1.2; Olympus Inc.). In FL mode, we scanned the sample stage to image raw stacks of Vph1-GFP-expressed yeasts with an axial spacing of 300 nm. These raw data were then deconvolved using the built-in software (Tomostudio, Tomocube Inc.) to generate processed volumetric images with enhanced resolution [21].

To acquire RI data, we captured 49 raw off-axis holograms in an off-axis Mach-Zehnder interferometer by angularly scanning a green plane wave using a digital micromirror device [22-24]. An RI tomogram was reconstructed from these holograms in the built-in software using the Rytov approximation and non-negativity constraints [25]. The theoretical lateral and axial resolution of HT were 110 nm and 360 nm, respectively [26]. The RI and FL data were registered and resized to achieve an isotropic voxel pitch of 200 nm.

### 2.3 Machine-learning-based segmentation

For high-throughput intracellular organization analysis, automated segmentation is key. We achieved this by implementing a machine-learning framework (Fig. 2). Initially, we utilized Ilastik, an open-source machine-learning annotation software, to prepare 3D labels of vacuoles [27] (Fig. 2a).

**Fig. 2.**
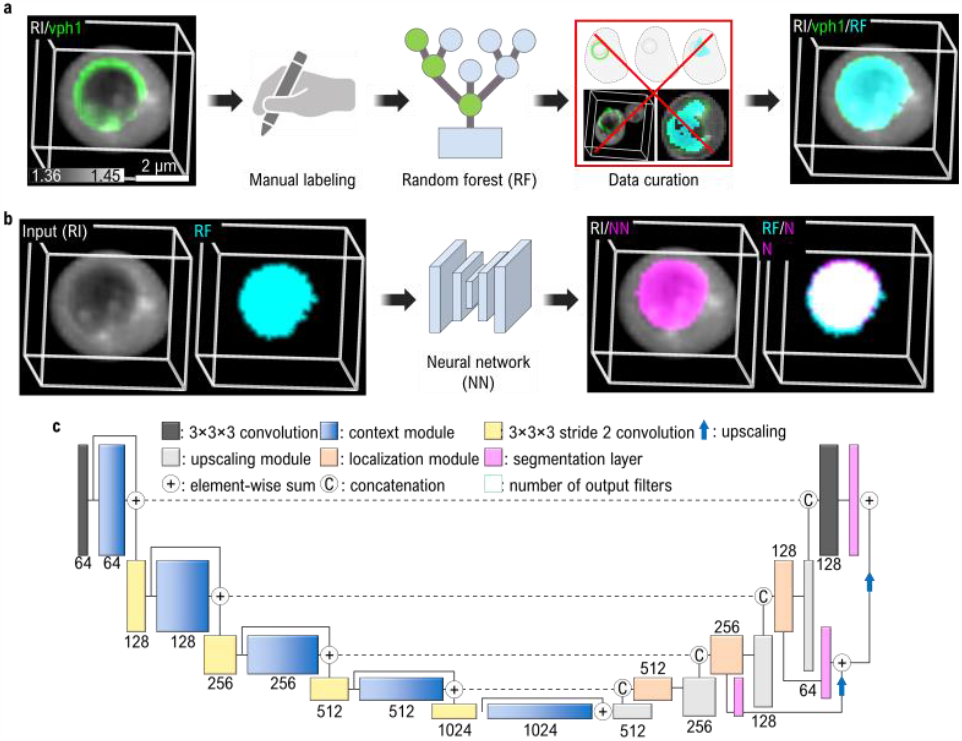
Data flowchart for automated 3D vacuole segmentation. (a) Sparse masks are annotated across XY, XZ, and YZ cross sections of 3D RI and FL Vph1-GFP images to prepare vacuole labels. These annotations are used to train a random forest segmentation algorithm, producing random-forest labels (RF). The dataset is then curated by excluding misregistered and poorly segmented data. (b) The neural network (NN) is trained using RI and the random-forest labels from (a) as inputs and training labels, respectively. The trained NN subsequently predicts 3D vacuole masks using only RI tomograms. (c) Depicts the architecture of the NN used for the segmentation task.

In each dataset, we manually annotated cytoplasm and vacuoles in at least 3 slices of XY, XZ, and YZ cross sections. A total of 163 individual cells from 21 randomly selected FL and RI datasets were annotated manually, training a random-forest classifier [28] built into the software. The following selected features were used; five Gaussian smoothing parameters (σ = 0.3, 0.7, 1.6, 3.5, and 10), three edge parameters (σ = 0.7, 1, and 5), and three texture parameters (σ = 1.6, 5, and 10) for each RI and FL image. This algorithm was then applied to 156 datasets, from which 69 datasets were curated as well-segmented, excluding those with artifacts due to movement during imaging or inaccurate segmentation.

To improve our framework and eliminate reliance on FL data, we introduced a deep NN designed to segment vacuoles using solely RI data (Fig. 2b). To train the network, we used the curated set of cells that were segmented based on the Vph1 vacuolar signal. The input images for this network were RI tomograms, normalized from [1.35, 1.45] float to [0, 255] integer formats. The labels used in the network were the same as those obtained from Ilastik.

Our network was adapted from a classical 3D U-Net used for brain tumor segmentation [28] (Fig. 2c). The structure comprises a down-sampling encoder pathway and up-sampling decoder pathway [29], each with its unique components. Each encoder path includes a context module that consists of two 3 × 3 × 3 convolution layers and a dropout layer (*p* = 0.75) in between. The context module is connected by a 3 × 3 × 3 convolution layer with an input stride 2. Each decoder path includes a localization module that has a 3 × 3 × 3 convolution layer followed by a layer that reduces the number of feature maps by half. The localization module is up-sampled and concatenated with the features from the corresponding level of the context module. A segmentation layer is set up at different levels, and the network outputs are combined to form the final network output. A segmentation layer is set up at different levels of the network, and the deep-learning outputs are combined via elementwise summation to form the final network output. Nonlinear activations are achieved through leaky ReLU activation functions with a 10^−2^ negative slope and instance normalization [30].

We trained the architecture on a cloud computing platform (Google Colaboratory Pro plus, Google Inc.) using Python Pytorch code. We segmented 354 individual cells from 69 datasets and divided them into 266 training and 88 validation sets. With a batch size of 8, the network was trained for 200 epochs. The loss function comprised 90% binary cross entropy with a logistic function and 10% focal Tversky function (α = 0.25, β = 0.75, and γ = 2) [31], minimized by the Adam optimizer [32] with a learning rate of 10^−4^.

### 2.4 Quantitative analysis of vacuole and cytoplasm

To obtain the segmented vacuole masks, we used either the random forest algorithm or the NN, depending on which algorithm provided larger Pearson coefficients with the RI image. For cytoplasm, we did not apply the binary erosion because conventional method based on thresholding was sufficient. In detail, we defined the cytoplasm mask by thresholding the RI with the average value between the mean cytoplasm RI defined in Ilastik and the mean vacuole RI, followed by binary erosion with one voxel. Note that we didn’t use the binary erosion and the RI average thresholding on the vacuole segmentation. We then converted the obtained RI to dry-mass concentration using the RI increment of 0.1907 mL/g [33].

## 3. Results and discussion

### 3.1 Validation of segmentation performance

We evaluated the segmentation performance of both the random forest classifier and the deep NN for vacuoles (Fig. 3). We first examined the cross-sections of individual cell data for both the training and untrained datasets (Figs. 3a, b). In the training dataset, we curated data of 211 cells for which either random-forest labels or deep-learning outputs were well-segmented (Figs. 3a and c). As expected, the comparison of RI, FL, random-forest labels (RF), and neural-network outputs indicated that both algorithms provided well-segmented vacuole masks. Additionally, we curated 268 cell data from the untrained dataset (Figs. 3b and d). We found that the random-forest labels were inaccurate when FL and RI images were misregistered or the cell moved during the long fluorescence imaging (see Fig. 3b, bottom). This issue is overcome by the NN, which robustly provided vacuole masks where the RI values were smaller than the other regions.

**Fig. 3.**
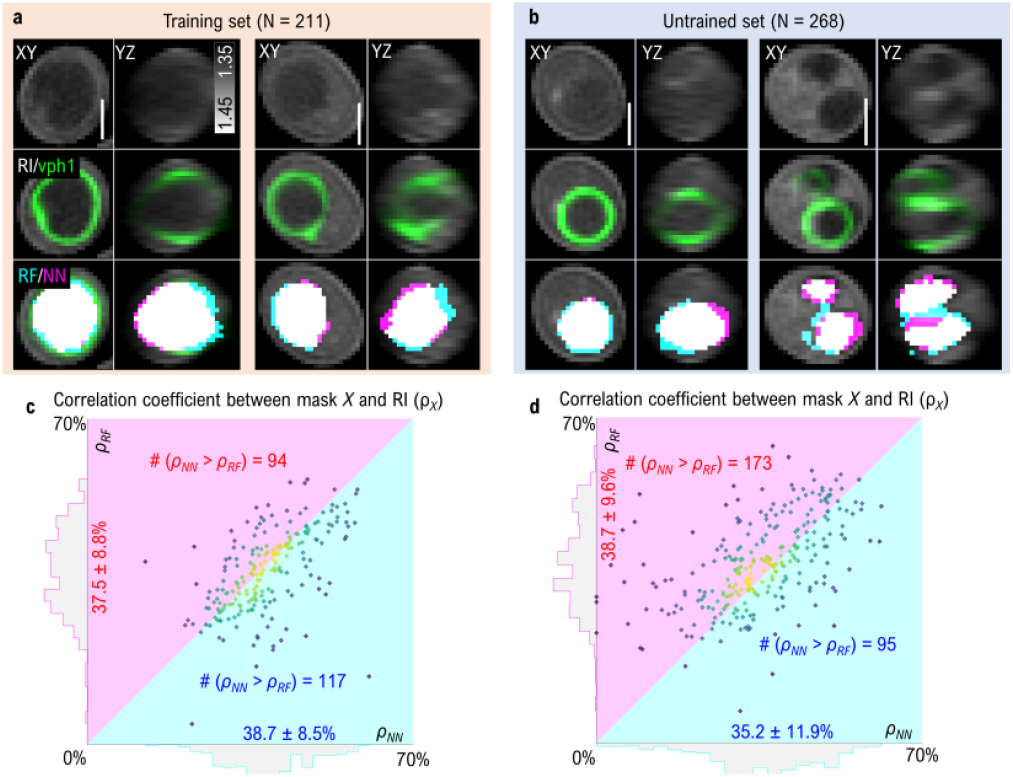
Segmentation performance evaluation. (a, b) Representative cross-sections of RI (gray), Vph1-GFP (green), label from random forest (cyan), and output from NN (magenta) for (a) the training dataset and (b) the untrained dataset, respectively. (c, d). Heat scatter maps and projected histograms of Pearson correlation coefficients (*ρ*) between 3D RI distributions and 3D cell masks for (c) the training dataset and (d) the untrained dataset, respectively. Axis label: (Horizontal, Vertical) = (*ρ*_output_, *ρ*_label_): The data are presented as mean ± standard deviation (SD). The magenta region indicates where the output mask from the NN has a higher *ρ* than the label from the random forest, and the cyan vice versa. Scale bar = 2 μm.

To quantify the segmentation performance of the algorithms, we computed the Pearson correlation coefficients between the obtained binary mask *X* and the RI image (*ρ*_*X*_). In both the training and untrained datasets, heat-scatter plots indicated a positive correlation between *ρ*_RF_ and *ρ*_*NN*_ (Figs. 3c, d). In the training set, the mean value of *ρ*_*RF*_ (mean ± SD = 38.7 ± 8.5%) was slightly larger than *ρ*_*NN*_ (mean ± SD = 37.5 ± 8.8%) (Fig. 3c). The number of datapoints with larger *ρ*_*RF*_ (magenta region in Fig. 3c) and larger *ρ*_*NN*_ (cyan region in Fig. 3c) were 117 and 94, respectively, which was expected because the NN was trained using the random-forest labels. In contrast, in the untrained dataset, the output masks from the NN output exhibited slightly larger correlation coefficients (*ρ*_*NN*_ = 38.7 ± 9.6%) than the random-forest labels with smaller standard deviations (*ρ*_*RF*_ = 35.2 ± 11.9%) (Fig. 3d). Correspondingly, the number of datapoints with larger *ρ*_*RF*_ (magenta region in Fig. 3d) and larger *ρ*_*NN*_ (cyan region in Fig. 3d) were 95 and 173, respectively.

Considering the visualized images and the quantitative results, both the random forest classifier and NN proved effective in segmenting 3D vacuole structures from sparse annotations. Crucially, deep learning provided prediction performances for 3D vacuoles comparable to the random forest algorithm without the need for FL images. Since deep learning is not affected by image registration artifacts caused by vacuole movement between the acquisition of the RI and FL images, it demonstrated slightly improved performance for untrained datasets. Collectively, the results suggest the viability of using deep learning for a more generalized analysis utilizing label-free RI images.

### 3.2 Quantitative analysis of cytoplasm and vacuoles

We utilized the established framework to perform quantitative biological analysis of cytoplasm and vacuoles in budding yeast cells under different conditions (Fig. 4). Budding yeast is a commonly used model organism for studying stress response effects, cell morphology and growth [34]. We investigated four conditions commonly used in yeast cell biology for which pre-existing knowledge on cytoplasmic properties is available. These conditions consist of exponentially growing cells (log phase, N = 96), nutrient depleted cells, (stationary phase, N = 141) and cells treated with the TORC1 inhibitor rapamycin in two conventional concentrations (0.1 μM (*N* = 82) and 1 μM (*N* = 92)).

**Fig. 4.**
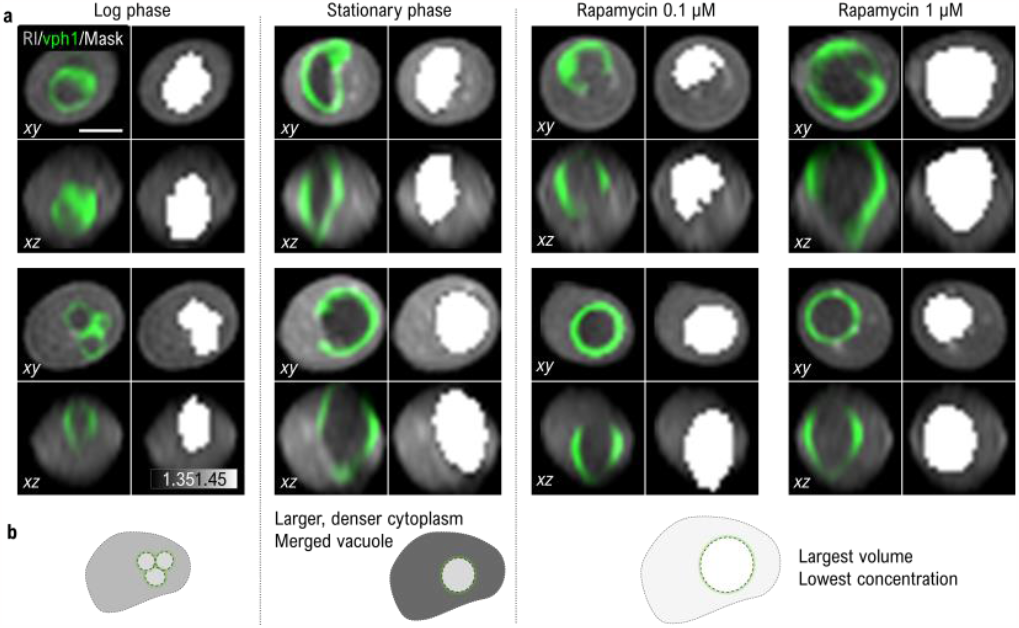
Biological application and quantitative analysis of budding yeast cells. (a) Representative cross-sections of RI (gray), FL (green; Vph1-GFP), and defined label (white) for four experimental groups: log phase, stationary phase, rapamycin 0.1 μM, and rapamycin 1 μM. (b) Schematic illustrating the overall results. Budding yeasts in log phase are used as controls. In the stationary phase, yeasts exhibit larger and denser cytoplasm. When the cells are treated with rapamycin, their volumes and dry masses increase while their concentration decreases. Scale bar = 2 μm.

Cells in stationary phase have been shown to be more refractile than log-phase cells [35]. Treatment with rapamycin was observed to increase cell volume and diffusion in the cytoplasm [9], which is an indicator for lower crowding and therefore lower dry-mass concentration. These known behaviors are reflected in our results (Fig. 4). Consistent with a previous study [34], the mean cell volumes in the log and stationary phases were 21.7 and 26.1 fL, respectively. The volume of the cytoplasm showed a significant increase in the stationary phase (Fig. 4c). Treatment with 1 μM rapamycin further increased the cytoplasmic volume, which is consistent with previous studies by Chan and Marshall [36].

The vacuole volume exhibited a similar trend except for the stationary phase. During the stationary phases, the size and mass of the cytoplasm increase, while the vacuoles retain their original size. In contrast, during the logarithmic phase, the vacuoles were fragmented, but under other conditions, they merge into one large vacuole (Fig. 4a). This is a well-known behavior [17, 37] that we addressed by adding up the separate vacuoles per cell for our analysis. However, small errors in segmentation can accumulate when comparing multiple vacuoles to one. Therefore, it is preferable to make comparisons between conditions that have only one vacuole, such as the stationary phase and rapamycin-treated conditions. In these conditions, the vacuole volume exhibited a significant, dose-dependent increase when treated with rapamycin, consistent with previous observations [33].

While the volume statistics (Fig. 5a) exhibited a similar trend to the dry mass (Fig. 5b), the median dry-mass concentrations showed stark differences (Fig. 5c). In the cytoplasm, the dry-mass concentration was largest in the stationary phase and decreased significantly when treated with increasing rapamycin concentration. This is in line with previous observations of increased crowding in starved cells [38] and decreased crowding upon rapamycin treatment [9]. In the vacuole, the dry-mass concentration remained comparable across conditions, except for a decrease in the 1 μM rapamycin treatment. All the above results follow previously observed patterns and therefore indicate that our method is robust to and able to quantify changes in both the cytoplasm and vacuole in different conditions.

**Fig. 5.**
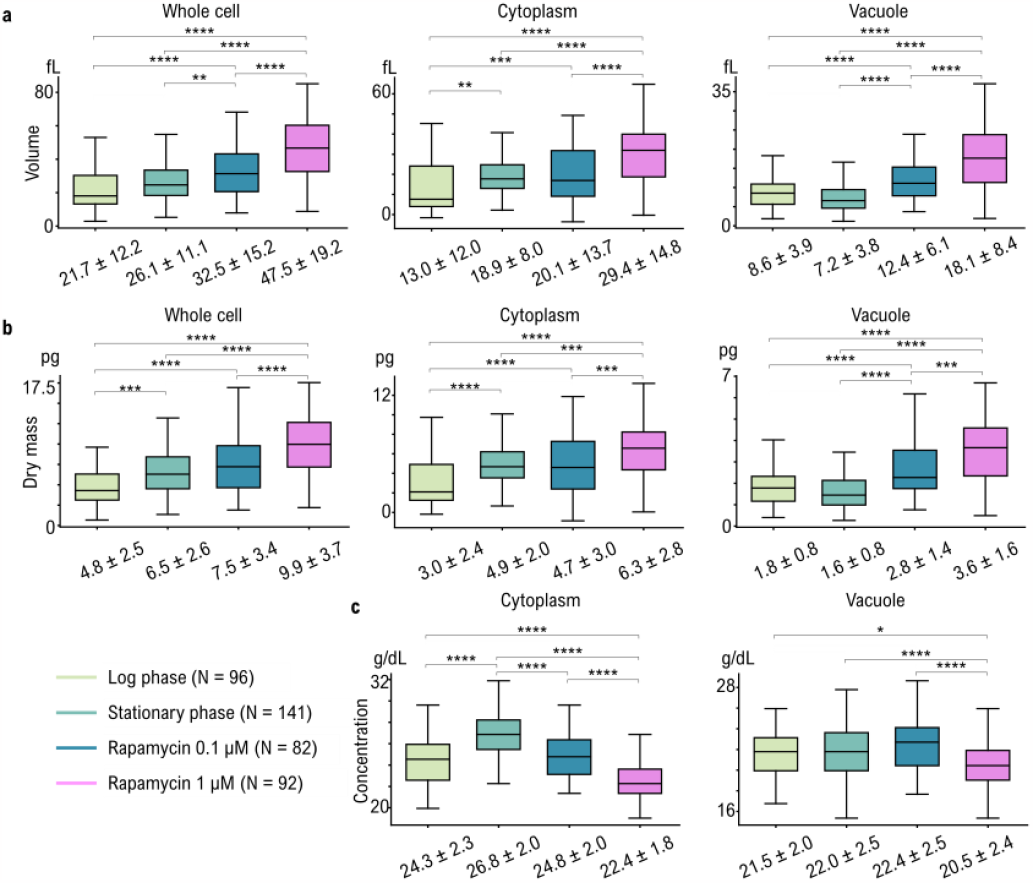
Biological application and quantitative analysis of budding yeast cells. (a) Quantitative results of volumetric parameters, including total cell volume, cytoplasm volume, and vacuole volume. (b) Quantitative results of dry mass parameters, including total cell dry mass, cytoplasm dry mass, and vacuole dry mass. (c) Quantitative results of median dry-mass concentration in cytoplasm and vacuoles. Tukey’s range test was performed, and the data are presented as mean ± standard deviation (SD). *: P<0.05, **: P<0.01, ***: P<0.001, ****: P<0.0001.

**Fig. 6.**
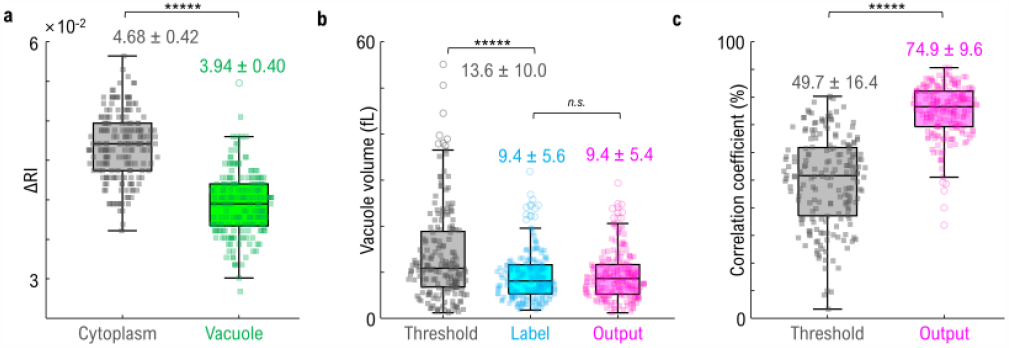
Comparison with conventional thresholding algorithm. (a) Statistical analysis of the relative refractive index to the background medium for defined cytoplasm (gray) and vacuole (green). (b) Vacuole volumes obtained from thresholding (gray; threshold) by the average value of the cytoplasm and the vacuole (ΔRI_thresh_= 4.31) and the random forest (cyan; label) and the deep-learning algorithm (magenta; output). (c) Pearson correlation coefficients of either the thresholding volume (gray) and the output from the deep NN (magenta) with the label from the random forest classifier. *n* = 192. Two-tailed paired Student t tests were performed. and the data are presented as mean ± standard deviation (SD). *****: *P* < 1^-10^. n.s.: nonsignificant *p* value.

## 4. Discussion & Conclusion

Our study presents a quantitative evaluation of intracellular components, achieved by combining RI tomography with a machine-learning-based algorithm. The absence of chemical specificity in RI was counterbalanced by correlative fluorescence imaging, enabling precise annotations of yeast vacuole. Even a minimal set of annotations was enough to kick-start the supervised volumetric segmentation of intracellular organization, facilitating high-throughput statistical analysis of budding yeasts.

Our approach bears similarities to previous studies employing machine learning for segmenting immunological synapses [39, 40] or predicting fluorescence images from RI [41]. However, our study stands apart in its data preparation stage, where we utilized Ilastik to significantly streamline the labor-intensive annotation procedure. This streamlining of dataset preparation is essential for generalizing machine-learning-based analysis tasks and enhancing accessibility.

Our label-free segmentation strategy succeeded fundamentally because there is a significant RI difference between the cytoplasm and the vacuole (Fig. 5a). One may then question whether a simple segmentation using a RI-thresholding algorithm should also work. As in Fig. 4e, however, this is not simple because the intracellular RI difference depends significantly upon the treatment and culturing conditions. We quantitatively validated this by comparing the volume obtained from a conventional RI-thresholding algorithm and our machine-learning algorithms (Figs. 5b, c). For paired Student *t* tests, we used the dataset where the vacuole masks obtained from both the random forest classifier and the deep NN were successfully segmented. The results indicated that the simple thresholding algorithm segmented larger vacuole volumes than the other algorithms (Fig. 5b). The similarity with the labels obtained from the random forest classifier was also compared by computing the Pearson correlation coefficients, which indicated significantly lower correlation when an RI-thresholding algorithm was used (Fig. 5c). Taken together, both RI and complex morphological features should be considered for accurate segmentation of objects in cell.

The segmentation performance can be further improved simply by updating the algorithms used in both Ilastik and our NN. It is important to note that the architecture used in our study is a basic model. We anticipate that future studies will enhance model performance by adopting recently developed algorithms such as diffusion models [42] or attention models [43, 44]. This will improve the accuracy and precision of segmentation, allowing for distinction of the vacuolar membrane and individual compartments of vacuoles in future studies.

The biological results presented here corroborate interesting correlations between cellular metabolic activities and the density of intracellular compartments. Unlike previous studies that focused on size and density of the entire cell, our method allows for simultaneous and detailed quantification of cytoplasmic and vacuolar densities, offering new possibilities for applications in yeast cell biology. This will facilitate research on regulatory mechanisms underlying cytoplasmic density as well as its effects on other processes, such as phase separation and pathologies. Additionally, combining our method with a microfluidic chip would allow for quantitative time-lapse measurements of the cell’s response to fast changes, such as glucose starvation, osmotic shock or temperature changes, which have all been implicated in affecting cytoplasmic density [14, 38, 45]. Moreover, it would allow us to follow cells over longer time-scales and track the effect of aging on cytoplasmic density [5]. Our method is available online (see Data availability) and we hope that it will prove a useful tool for the analysis of label free images.

## Funding

This work was supported by the grant of Young researchers’ exchange programme between Korea and Switzerland (KR_EG_042022_05) from ETH Zurich as the Leading House Asia, National Research Foundation of Korea (2015R1A3A2066550, 2022M3H4A1A02074314), Swiss National Science Foundation (SNSF) Eccellenza (Nr. PCEFP3 187003), Institute of Information & communications Technology Planning & Evaluation (IITP; 2021-0-00745) grant funded by the Korea government (MSIT), and KAIST Institute of Technology Value Creation, Industry Liaison Center (G-CORE Project) grant funded by MSIT (N11230131), Tomocube Inc..

## Acknowledgments

We thank Dr. Geon Kim, Mr. Dohyeon Lee, and Ms. Juyeon Park in Biomedical Optics Laboratory, KAIST, for helpful discussion on machine learning.

## Disclosures

ML and YKP have financial interests in Tomocube Inc., a company that commercializes HT and is one of the sponsors of the work. The other authors declare no conflicts of interest.

## Data availability

The used dataset and codes are available in https://drive.google.com/drive/folders/1AS_LFlA2SfKYHoJge6VNloMOnMaNewoK?usp=share_link

## References

1. M. Schürmann, J. Scholze, P. Müller, J. Guck, and C. J. Chan, “Cell nuclei have lower refractive index and mass density than cytoplasm,” Journal of biophotonics 9, 1068–1076 (2016).

2. S. Abuhattum, K. Kim, T. M. Franzmann, A. Eßlinger, D. Midtvedt, R. Schlüßler, S. Möllmert, H.-S. Kuan, S. Alberti, and V. Zaburdaev, “Intracellular mass density increase is accompanying but not sufficient for stiffening and growth arrest of yeast cells,” Frontiers in Physics 6, 131 (2018).

3. W. H. Grover, A. K. Bryan, M. Diez-Silva, S. Suresh, J. M. Higgins, and S. R. Manalis, “Measuring single-cell density,” Proceedings of the National Academy of Sciences 108, 10992–10996 (2011).

4. B. R. Parry, I. V. Surovtsev, M. T. Cabeen, C. S. O’Hern, E. R. Dufresne, and C. Jacobs-Wagner, “The bacterial cytoplasm has glass-like properties and is fluidized by metabolic activity,” Cell 156, 183–194 (2014).

5. G. E. Neurohr, R. L. Terry, J. Lengefeld, M. Bonney, G. P. Brittingham, F. Moretto, T. P. Miettinen, L. P. Vaites, L. M. Soares, and J. A. Paulo, “Excessive cell growth causes cytoplasm dilution and contributes to senescence,” cell 176, 1083–1097. e1018 (2019).

6. V. C. Hecht, L. B. Sullivan, R. J. Kimmerling, D.-H. Kim, A. M. Hosios, M. A. Stockslager, M. M. Stevens, J. H. Kang, D. Wirtz, and M. G. Vander Heiden, “Biophysical changes reduce energetic demand in growth factor–deprived lymphocytes,” Journal of Cell Biology 212, 439–447 (2016).

7. K. L. Cooper, S. Oh, Y. Sung, R. R. Dasari, M. W. Kirschner, and C. J. Tabin, “Multiple phases of chondrocyte enlargement underlie differences in skeletal proportions,” Nature 495, 375–378 (2013).

8. S. Oh, C. Lee, W. Yang, A. Li, A. Mukherjee, M. Basan, C. Ran, W. Yin, C. J. Tabin, and D. Fu, “Protein and lipid mass concentration measurement in tissues by stimulated Raman scattering microscopy,” Proceedings of the National Academy of Sciences 119, e2117938119 (2022).

9. M. Delarue, G. P. Brittingham, S. Pfeffer, I. Surovtsev, S. Pinglay, K. Kennedy, M. Schaffer, J. Gutierrez, D. Sang, and G. Poterewicz, “mTORC1 controls phase separation and the biophysical properties of the cytoplasm by tuning crowding,” Cell 174, 338–349. e320 (2018).

10. Y. Shin, and C. P. Brangwynne, “Liquid phase condensation in cell physiology and disease,” Science 357, eaaf4382 (2017).

11. T. Kim, J. Yoo, S. Do, D. S. Hwang, Y. Park, and Y. Shin, “RNA-mediated demixing transition of low-density condensates,” Nature Communications 14, 2425 (2023).

12. R. Babazadeh, C. B. Adiels, M. Smedh, E. Petelenz-Kurdziel, M. Goksör, and S. Hohmann, “Osmostress-induced cell volume loss delays yeast Hog1 signaling by limiting diffusion processes and by Hog1-specific effects,” PLoS One 8, e80901 (2013).

13. S. Krishnaswamy, M. H. Spitzer, M. Mingueneau, S. C. Bendall, O. Litvin, E. Stone, D. Pe’er, and G. P. Nolan, “Conditional density-based analysis of T cell signaling in single-cell data,” Science 346, 1250689 (2014).

14. B. D. Knapp, P. Odermatt, E. R. Rojas, W. Cheng, X. He, K. C. Huang, and F. Chang, “Decoupling of rates of protein synthesis from cell expansion leads to supergrowth,” Cell systems 9, 434–445. e436 (2019).

15. M. Mir, Z. Wang, Z. Shen, M. Bednarz, R. Bashir, I. Golding, S. G. Prasanth, and G. Popescu, “Optical measurement of cycle-dependent cell growth,” Proceedings of the National Academy of Sciences 108, 13124–13129 (2011).

16. T. A. Zangle, and M. A. Teitell, “Live-cell mass profiling: an emerging approach in quantitative biophysics,” Nature methods 11, 1221–1228 (2014).

17. L. Wang, E. S. Seeley, W. Wickner, and A. J. Merz, “Vacuole fusion at a ring of vertex docking sites leaves membrane fragments within the organelle,” Cell 108, 357–369 (2002).

18. Y. Park, C. Depeursinge, and G. Popescu, “Quantitative phase imaging in biomedicine,” Nature photonics 12, 578–589 (2018).

19. R. Barer, K. Ross, and S. Tkaczyk, “Refractometry of living cells,” Nature 171, 720–724 (1953).

20. K. Kim, W. S. Park, S. Na, S. Kim, T. Kim, W. Do Heo, and Y. Park, “Correlative three-dimensional fluorescence and refractive index tomography: bridging the gap between molecular specificity and quantitative bioimaging,” Biomedical Optics Express 8, 5688–5697 (2017).

21. D. S. Biggs, and M. Andrews, “Acceleration of iterative image restoration algorithms,” Applied optics 36, 1766–1775 (1997).

22. S. Shin, K. Kim, J. Yoon, and Y. Park, “Active illumination using a digital micromirror device for quantitative phase imaging,” Opt Lett 40, 5407–5410 (2015).

23. K. Lee, K. Kim, G. Kim, S. Shin, and Y. Park, “Time-multiplexed structured illumination using a DMD for optical diffraction tomography,” Opt Lett 42, 999–1002 (2017).

24. S. Shin, K. Kim, T. Kim, J. Yoon, K. Hong, J. Park, and Y. Park, “Optical diffraction tomography using a digital micromirror device for stable measurements of 4D refractive index tomography of cells,” in Quantitative Phase Imaging II(SPIE2016), pp. 156–163.

25. J. Lim, K. Lee, K. H. Jin, S. Shin, S. Lee, Y. Park, and J. C. Ye, “Comparative study of iterative reconstruction algorithms for missing cone problems in optical diffraction tomography,” Optics express 23, 16933–16948 (2015).

26. C. Park, S. Shin, and Y. Park, “Generalized quantification of three-dimensional resolution in optical diffraction tomography using the projection of maximal spatial bandwidths,” JOSA A 35, 1891–1898 (2018).

27. S. Berg, D. Kutra, T. Kroeger, C. N. Straehle, B. X. Kausler, C. Haubold, M. Schiegg, J. Ales, T. Beier, and M. Rudy, “Ilastik: interactive machine learning for (bio) image analysis,” Nature methods 16, 1226–1232 (2019).

28. F. Isensee, P. Kickingereder, W. Wick, M. Bendszus, and K. H. Maier-Hein, “Brain tumor segmentation and radiomics survival prediction: Contribution to the brats 2017 challenge,” in Brainlesion: Glioma, Multiple Sclerosis, Stroke and Traumatic Brain Injuries: Third International Workshop, BrainLes 2017, Held in Conjunction with MICCAI 2017, Quebec City, QC, Canada, September 14, 2017, Revised Selected Papers 3(Springer 2018), pp. 287–297.

29. Ö. Çiçek, A. Abdulkadir, S. S. Lienkamp, T. Brox, and O. Ronneberger, “3D U-Net: learning dense volumetric segmentation from sparse annotation,” in Medical Image Computing and Computer-Assisted Intervention–MICCAI 2016: 19th International Conference, Athens, Greece, October 17-21, 2016, Proceedings, Part II 19(Springer 2016), pp. 424–432.

30. D. Ulyanov, A. Vedaldi, and V. Lempitsky, “Instance normalization: The missing ingredient for fast stylization,” arXiv preprint arXiv:1607.08022 (2016).

31. N. Abraham, and N. M. Khan, “A novel focal tversky loss function with improved attention u-net for lesion segmentation,” in 2019 IEEE 16th international symposium on biomedical imaging (ISBI 2019)(IEEE 2019), pp. 683–687.

32. D. P. Kingma, and J. Ba, “Adam: A method for stochastic optimization,” arXiv preprint arXiv:1412.6980 (2014).

33. H. Zhao, P. H. Brown, and P. Schuck, “On the distribution of protein refractive index increments,” Biophysical journal 100, 2309–2317 (2011).

34. D. Midtvedt, E. Olsén, F. Höök, and G. D. Jeffries, “Label-free spatio-temporal monitoring of cytosolic mass, osmolarity, and volume in living cells,” Nature Communications 10, 340 (2019).

35. V. Bugeja, J. Piggott, and B. Carter, “Differentiation of Saccharomyces cerevisiae at slow growth rates in glucose-limited chemostat culture,” Microbiology 128, 2707–2714 (1982).

36. Y.-H. M. Chan, and W. F. Marshall, “Organelle size scaling of the budding yeast vacuole is tuned by membrane trafficking rates,” Biophysical journal 106, 1986–1996 (2014).

37. S. P. Rayermann, G. E. Rayermann, C. E. Cornell, A. J. Merz, and S. L. Keller, “Hallmarks of reversible separation of living, unperturbed cell membranes into two liquid phases,” Biophysical journal 113, 2425–2432 (2017).

38. R. P. Joyner, J. H. Tang, J. Helenius, E. Dultz, C. Brune, L. J. Holt, S. Huet, D. J. Müller, and K. Weis, “A glucose-starvation response regulates the diffusion of macromolecules,” elife 5, e09376 (2016).

39. M. Lee, Y.-H. Lee, J. Song, G. Kim, Y. Jo, H. Min, C. H. Kim, and Y. J. E. Park, “Deep-learning-based three-dimensional label-free tracking and analysis of immunological synapses of CAR-T cells,” eLife 9, e49023 (2020).

40. J. Choi, H.-J. Kim, G. Sim, S. Lee, W. S. Park, J. H. Park, H.-Y. Kang, M. Lee, W. D. Heo, and J. Choo, “Label-free three-dimensional analyses of live cells with deep-learning-based segmentation exploiting refractive index distributions,” bioRxiv, 2021.2005. 2023.445351 (2021).

41. Y. Jo, H. Cho, W. S. Park, G. Kim, D. Ryu, Y. S. Kim, M. Lee, S. Park, M. J. Lee, and H. Joo, “Label-free multiplexed microtomography of endogenous subcellular dynamics using generalizable deep learning,” Nature Cell Biology 23, 1329–1337 (2021).

42. J. Sohl-Dickstein, E. Weiss, N. Maheswaranathan, and S. Ganguli, “Deep unsupervised learning using nonequilibrium thermodynamics,” in International Conference on Machine Learning(PMLR 2015), pp. 2256–2265.

43. A. Vaswani, N. Shazeer, N. Parmar, J. Uszkoreit, L. Jones, A. N. Gomez, Ł. Kaiser, and I. Polosukhin, “Attention is all you need,” Advances in neural information processing systems 30 (2017).

44. J. Fu, J. Liu, H. Tian, Y. Li, Y. Bao, Z. Fang, and H. Lu, “Dual attention network for scene segmentation,” in Proceedings of the IEEE/CVF conference on computer vision and pattern recognition(2019), pp. 3146–3154.

45. L. B. Persson, V. S. Ambati, and O. Brandman, “Cellular control of viscosity counters changes in temperature and energy availability,” Cell 183, 1572–1585. e1516 (2020).

